# A Novel Humanized Immune Stroma PDX Cancer Model for Therapeutic Studies

**DOI:** 10.1101/2023.07.03.547206

**Authors:** Dongli Yang, Ian Beddows, Huijuan Tang, Sandra Cascio, Stacy C. McGonigal, Shoumei Bai, Benjamin K. Johnson, John J. Powers, Rajesh Acharya, Riyue Bao, Tullia C. Bruno, Thing R. Soong, Jose R. Conejo-Garcia, Hui Shen, Moses T. Bility, Ronald J. Buckanovich

## Abstract

Standard preclinical human tumor models lack a human tumor stroma. However, as stroma contributes to therapeutic resistance, the lack of human stroma may make current models less stringent for testing new therapies. To address this, using patient-derived tumor cells, patient derived cancer-associated mesenchymal stem/progenitor cells, and human endothelial cells, we created a Human Stroma-Patient Derived Xenograft (HS-PDX) tumor model. HS-PDX, compared to the standard PDX model, demonstrate greater resistance to targeted therapy and chemotherapy, and better reflect patient response to therapy. Furthermore, HS-PDX can be grown in mice with humanized bone marrow to create humanized immune stroma patient-derived xenograft (HIS-PDX) models. The HIS-PDX model contains human connective tissues, vascular and immune cell infiltrates. RNA sequencing analysis demonstrated a 94-96% correlation with primary human tumor. Using this model, we demonstrate the impact of human tumor stroma on response to CAR-T cell therapy and immune checkpoint inhibitor therapy. We show an immunosuppressive role for human tumor stroma and that this model can be used to identify immunotherapeutic combinations to overcome stromally mediated immunosuppression. Combined, our data confirm a critical role for human stoma in therapeutic response and indicate that HIS-PDX can be an important tool for preclinical drug testing.

**Statement of Significance:** We developed a tumor model with human stromal, vascular, and immune cells. This model mirrors patient response to chemotherapy, targeted therapy, and immunotherapy, and can be used to study therapy resistance.

## Introduction

While initial studies of therapeutic resistance were focused on the tumor cells, the past decade of research has clearly demonstrated that non-tumor host cells in the tumor microenvironment (TME) play a crucial role in therapeutic resistance. Cancer-associated fibroblasts (CAF), mesenchymal stem/progenitor cells (MSC), vascular endothelial cells (EC) and immune cells (IC) can all impact therapeutic response (1,2). The biophysical aspects of tumor stroma can alter drug diffusion (3) and directly impact the cancer cell to promote therapeutic resistance to multiple agents (4). Host cell production of growth factors and chemokines can also induce therapeutic resistance in cancer cells (5). Host cells also critically regulate anti-tumor immunity. Tumor vasculature can prevent effector cells from infiltrating tumor islets(6). Similarly, MSC are linked with effector immune cell exclusion and recruitment of immunosuppressive myeloid cells which can render effector cells inactive (7,8).

Given the critical role of stromal cells in tumor therapeutic response, presence of human tumor stroma in preclinical drug testing models is essential. Unfortunately, cell line xenografts, which are the most used human tumor models, completely lack human stroma. Similarly, while early passage patient-derived xenografts (PDX) may contain stromal cells and be more predictive of outcome, PDX need to be expanded for drug testing and with passage human stroma is replaced by mouse stroma (9,10). More recently, numerous ‘humanized’ PDX models, incorporating human immune cells have been developed (11–13). These models have many strengths, but often have an underrepresentation of the myeloid cell compartment and lack other human stromal components such as human fibroblast and vascular cells. Thus, current humanized PDX models, while being a clear improvement over standard xenografts, do not completely reflect the human TME.

Deficiencies in preclinical models used in drug development can have significant repercussions. Poor models of cancer used in drug development are proposed to play a major role in the ultimate failure of drugs in clinical trials (14). Drugs which show significant activity in mouse models of cancer, are often ineffective in patients in clinical trials(15). In addition to increasing the likelihood that patients enrolled in clinical trials will not experience clinical benefit, these failures come with substantial financial costs which increase the cost of drug development and lead to high drug costs for patients.

To overcome the deficiencies in the current humanized mouse cancer models, we co-engrafted primary human tumor cells with multipotent cancer-associated MSC (CA-MSC -- which can differentiate into fibroblasts, myofibroblasts, and adipocytes) and various endothelial sources to generate a humanized stroma-PDX (HS-PDX). HS-PDX were then grown in mice with human bone-marrow/liver/thymic+/-splenic (BLT/S) transplanted humanized immune systems, creating humanized immune stroma-PDX (HIS-PDX). Molecular and protein profiling indicate these tumors closely reflect primary human tumors. We find HS-PDX model compared to standard xenografts and PDX, like human disease, is more resistant to targeted therapies. Further we find the HIS-PDX model can be used to evaluate human immunotherapy approaches, and better understand the biology of human cancer.

## Results

### CA-MSC derived stroma impacts therapeutic response

We previously showed, using human cell line xenografts that CA-MSC differentiate to form a desmoplastic human stroma, including fibroblasts, myofibroblasts, and adipocytes (16,17). To determine if addition of human stroma could impact therapeutic response, we first used cell line xenograft models combined with CA-MSC as a source of human stroma. As the Notch inhibitors showed early promise in ovarian cancer preclinical studies, but ultimately failed in clinical trial, we initially tested dibenzazepine (DBZ), a gamma-secretase inhibitor targeting the Notch signaling pathway(18,19). While DBZ significantly reduced cell line xenograft growth in both SKOV3 and HEY1 cell line xenografts, there was no anti-tumor activity observed when tumor cells were xenografted with CA-MSC (Fig. 1a). We next tested therapeutic inhibitors of IL6 signaling. Like DBZ, anti-human IL6 neutralizing antibodies effectively restricted cell line xenograft growth in the absence of human stroma but were ineffective in the presence of human stroma. In contrast, the JAK-STAT inhibitor, TG101209, restricted tumor growth in both the absence and presence of human stroma (Fig. 1b). Suggesting an on-target effect, tumor response correlated with inhibition of pSTAT3 (Fig. 1c).

**Figure 1:**
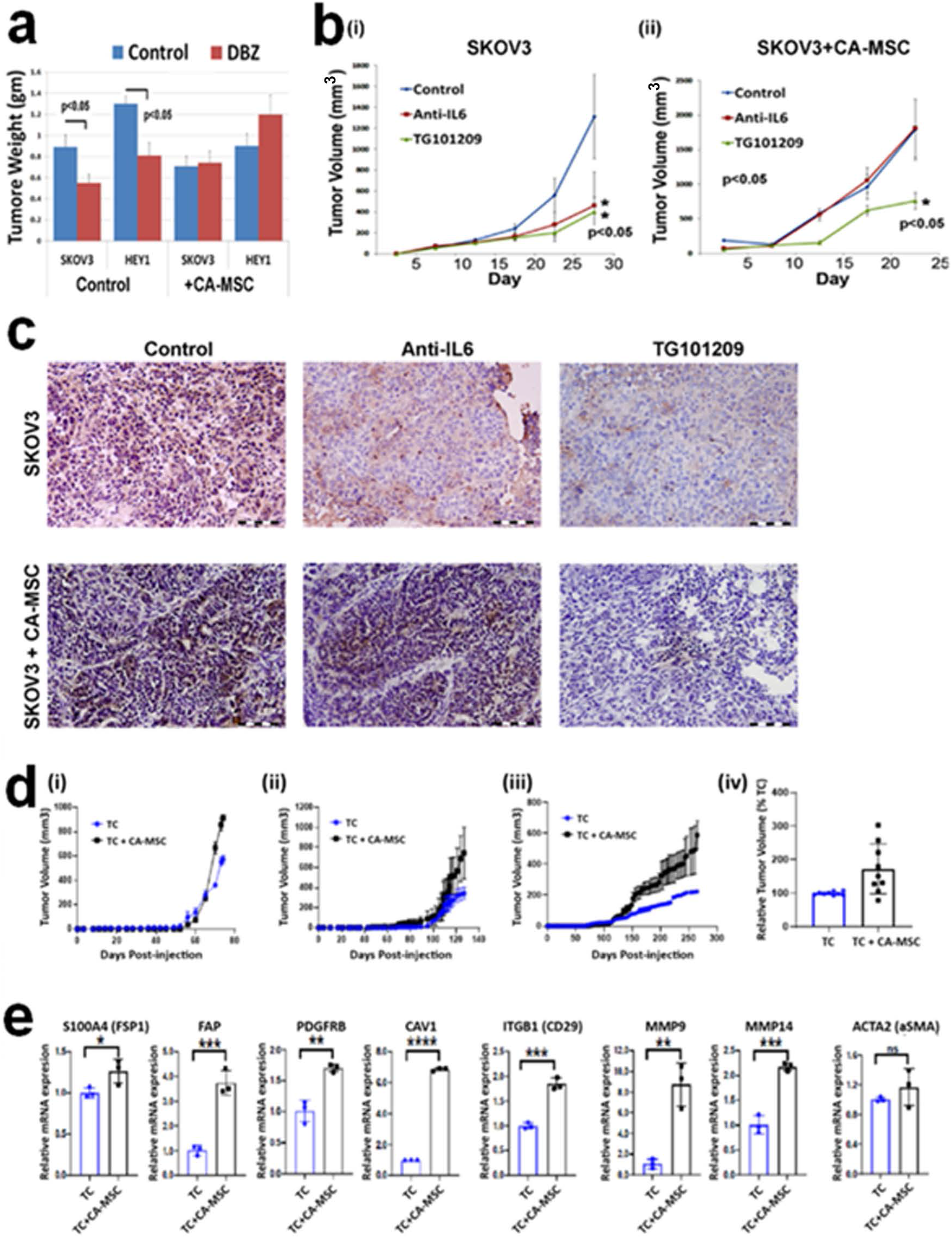
Effects of CA-MSC on the response of targeted therapy response and stromal gene expression. (**a**) Final tumor weights of control and dibenzazepine (DBZ) treated SKOV3 and HEY1 cell line xenografts (CDX) in the absence or presence of CA-MSC. (**b**) Tumor growth curves of cell line xenografts generated in the absence or presence of CA-MSC and treated with either anti-human IL6 neutralizing antibodies or JAK-STAT inhibitor (TG101209). (**c**) pSTAT3 IHC for the indicated tumor treatment groups. Scale bars, 200 µm. (**di-iii**) PDX Tumor growth curves for tumor cells grown alone or in the presence of CA-MSC from three different patients. (**div**) a box chart showing summary of relative tumor volumes at end of experiments in PDXs engrafted with TC alone (n = 9) or TC + CA-MSC (n = 9). Tumor volumes were normalized to the mean of tumor volumes in each Pt. PDX engrafted with TC alone. (**e**) qRT-PCR for human-specific stromal gene expression in PDX models with or without CA-MSC. GAPDH was used as an internal control. TC, NSG mice were engrafted with PDX TC alone; TC + CA-MSC, NSG mice were engrafted with PDX TC and CA-MSC. \**p* < 0.05, \*\**p* < 0.01, \*\*\**p* < 0.001, \*\*\*\**p* < 0.0001.

To move this study beyond cell line xenografts, we next combined human CA-MSC with primary human tumor samples to establish PDX with partially humanized stroma. Primary tumor cells were derived from either tumor-associated ascites or live single cell suspensions of primary tumors. Incorporation of CA-MSC increased tumor growth rates and significantly reduced time to tumor harvest (Fig. 1d). Real-time quantitative reverse transcription PCR (qRT-PCR) analysis confirmed that the incorporation of CA-MSC was associated with increased expression of numerous stromal mRNAs including fibroblast associated protein (FAP), S100A4, PDGFRβ, CAV1, ITGB1, MMP9 and MMP14 (Fig. 1e) (20–24). In addition, we observed that CA-MSC increase tumor metastasis to ovaries (Supplemental Fig. 1). This finding is consistent with our prior studies (25) and suggest this model reflects cancer biology better than cell line xenografts.

### Incorporating human vasculature to create a humanized stroma-PDX (HS-PDX)

While MSC have been reported to generate endothelial cells(26), we did not observe formation of human blood vessels with the incorporation of CA-MSC alone. To incorporate human tumor blood vessels in the HS-PDX, we tested different sources of endothelial cells (EC). As human endothelial progenitor cells (EPC) have been reported to form vessels in mice(27), EPC from two different vendors were tested. However, in our hands, EPC injection *in vivo* (grown alone or in combination with tumor cells/MSC xenografts) resulted in the formation of teratomas. Thus, these EPC sources were no longer studied. Next, we tested human induced pluripotent stem cells (iPSC)-derived EPC (iPSC-EPC) as described in Methods section and Supplemental Figure 2 legend. iPSC-EPC did not form tumors/teratomas (Supplemental Fig. 2b) and addition of iPSC-EPC to cell line xenografts generated clear human CD31^+^ vessels (Supplemental Fig. 2c). However, human vessels were relatively rare.

Human CD34^+^KDR^+^NRP1^+^ cells are also reported to form functional blood vessels in mice(28). We therefore tested the impact of patient tumor derived CD34^+^KDR^+^NRP1^+^ cells (Pt-EPC) FACS isolated from primary ovarian tumor specimens. CA-MSC and Pt-EPC grown in the absence of tumor cells demonstrated no growth (Fig. 2ai). The addition of Pt-EPC to tumor cells + CA-MSC significantly increased tumor growth (Fig. 2ai-ii). Histological analysis of tumors confirmed the formation of a robust human CD31^+^ tumor vasculature (Fig. 2b). Furthermore, these vessels also express the human ovarian tumor vascular specific marker EGFL6 (Fig. 2c) (29,30).

**Figure 2:**
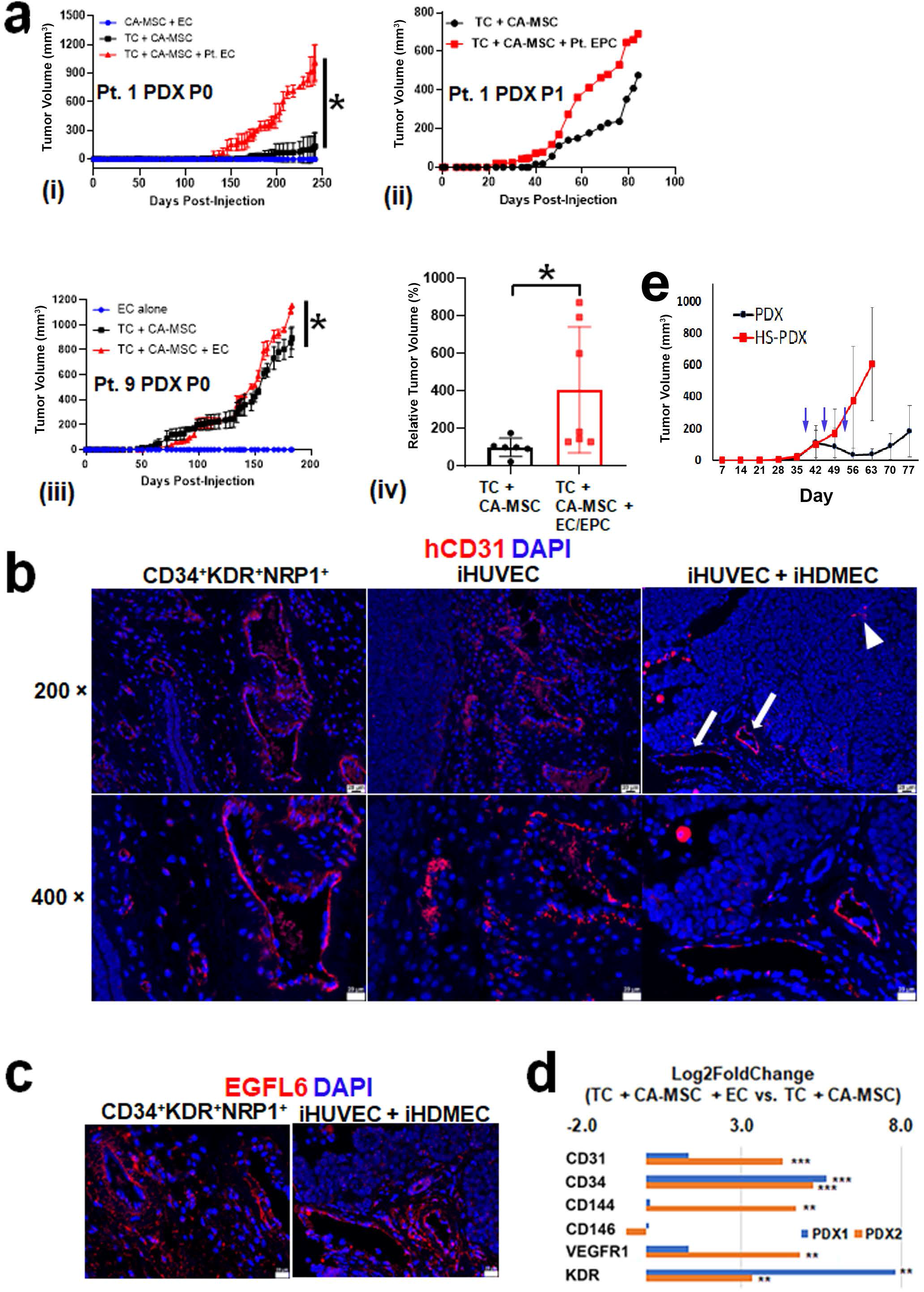
Generating a humanized stroma PDX (HS-PDX). (**a**) Tumor growth curves (i – iii) and (iv) summary of relative final tumor volumes of HS-PDXs with the indicate EC/EPC sources. TC + CA-MSC in the absence or presence of human EC (TC + CA-MSC; n = 6) (iHDMEC + iHUVEC) and Pt. EC/EPC (TC + CA-MSC + EC/EPC; n = 7). (**b**) IF evaluation of human tumor vascular antigen CD31 (red) in PDXs with the indicated EC/EPC sources. 4′,6-diamidino-2-phenylindole (DAPI; blue) is used to label cell nuclei. (**c**) IF evaluation of a tumor specific vascular marker EGFL6 (red). Cell nuclei were counterstained with DAPI (blue). (**d**) RNASeq based relative mRNA expression for two patient PDXs without and with the addition of human endothelial cells. **(e)** Growth curves for standard PDX and HS-PDX, derived from a patient with platinum refractory ovarian cancer, treated with carboplatin (blue arrows indicate timing of carboplatin administration. *Abbreviations*: CA-MSC, cancer-associated mesenchymal stem cells; EC, endothelial cells; EPC, endothelial progenitor cells; iHDMEC, immortalized human dermal microvascular endothelial cells; iHUVEC, immortalized human umbilical vein endothelial cells; PDX, patient-derived xenograft; P0 and P1, passage 0 and passage 1 (The first generation of PDX is denoted P0, and P0 is subsequently passaged to second generation (P1); Pt., patient; TC, tumor cells. \**p* < 0.05, \*\**p* < 0.01, \*\*\**p* < 0.001. Scale bars, 20 µm.

While Pt-EPC are a viable source of human tumor vasculature, Pt-EPC numbers are limited and thus difficult to use in large-scale therapeutic studies. In attempt to identify a more ‘user friendly’ source of human vasculature, we tested immortalized human umbilical vein endothelial cells (iHUVEC) and immortalized human dermal microvascular endothelial cells (iHDMEC). When iHUVEC added to TC + CA-MSC, tumors developed relatively large human CD31^+^ blood vessels located in the tumor periphery, but fewer microvessels (Fig. 2b). Combination of both iHUVEC and iHDMEC added to TC + CA-MSC, resulted in human CD31^+^ blood vessels located in both the peripheral (arrows) and central areas (arrowhead) of tumors (Fig. 2b). As with the tumor-derived EPC, iHUVEC/iHDMEC-derived human vessels also expressed human EGFL6 (Fig. 2c). Indeed, mRNA expression levels of numerous vascular antigens, such as CD31 and CD34, were also increased in PDXs generated by co-engraftment of TC + CA-MSC + EC (iHUVEC + iHDMEC) (Fig. 2d). Inclusion of human stroma also impacted tumor engraftment. Notably, the addition of both CA-MSC and EC to tumor cells increased successful PDX engraftment from 64.3% with traditional PDX to 81.8% (Supplemental Table 1). Combined these results suggest the addition of CA-MSC and immortalized endothelial cells can recreate a significant portion of the human stromal cell compartment and increase PDX growth and engraftment.

We defined the tumor model, which combines human patient tumor cells, CA-MSC, and endothelial cells, as a humanized stroma-PDX (HS-PDX). To evaluate the impact of HS-PDX on therapeutic response we tested the response of a PDX and HS-PDX derived from a patient with platinum and taxane refractory ovarian cancer. The standard PDX demonstrated an initial response to carboplatin and paclitaxel and then progressed with cessation of therapy. In contrast, the HS-PDX – mirroring the patient’s disease response– was completely refractory to carboplatin and paclitaxel therapy. This suggest the HS-PDX may better reflect patient response to therapy.

### Incorporating a Human Immune System to Generate Humanized Immune Stroma-PDX (HIS-PDX)

While the HS-PDX offers advantages compared to a standard PDX, it still lacks a human immune system. We therefore engrafted the HS-PDX model into several murine models with humanized bone marrow to identify the most robust Humanized Immune Stroma PDX (HIS-PDX). Specifically, we engrafted the HS-PDX model (patient tumor cells, CA-MSC, iHUVEC, and iHDMEC) into (i) human CD34^+^ hematopoietic stem cell-engrafted NSG mice (HuNSG-PDX), (ii) human CD34^+^ hematopoietic stem cell-engrafted into mice expressing human IL3, GM-CSF (CSF2) and SCF (KITLG) transgenic mice (HuSGM3-PDX), and (iii) NSG mice transplanted with human bone-marrow, liver, thymus (without or with spleen) (BLT(S)-PDX) mouse models (31). Resultant tumors were analyzed by RNA-seq, comparing the expression of immune cells in each tumor, normalized to expression in matched primary human tumor.

Consistent with prior studies, lymphocyte-associated genes (CD3, CD4, CD8, etc) were well represented in both the HuSGM3-HIS-PDX and BLT-HIS-PDX models (Fig. 3a). In contrast, myeloid cell markers (CD11b, CD14, CD80, CD163) (Fig. 3b) and natural killer (NK) cell markers (CD16, NKp46, GNLY, GZMB) (Fig. 3c) were relatively under-represented in the HuSGM3 mice while the BLT-PDX more reflected the primary tumor. Immunofluorescence (IF) confirmed the presence of tumor infiltrating CD3^+^ and CD8^+^ lymphocytes in both HuSGM3 and BLT mice, with T cells in BLT mice generally restricted to the peritumoral stroma (Fig. 3d). IF revealed CD14^+^ and CD163^+^ tumor-associated myeloid cells only in the BLT mice (Fig. 3d). Combined these studies suggest the BLT model has the strongest representation of human tumor infiltrating leukocytes, with the myeloid cell compartment in this model most reflecting that seen in primary human tumors.

**Figure 3:**
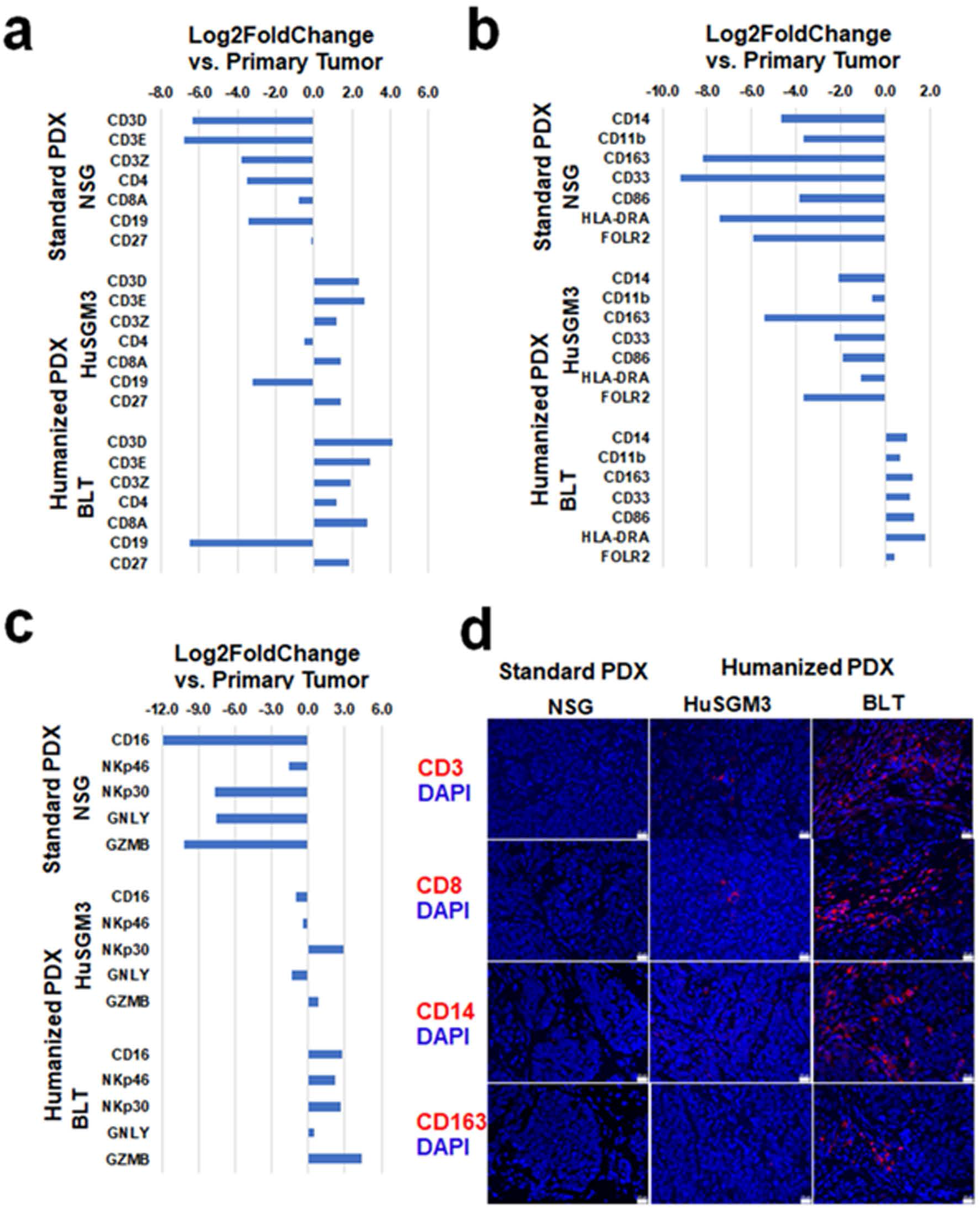
Identifying the optimal humanized bone marrow source for the humanized immune stroma PDX (HIS-PDX). PDX models established in immunodeficient NSG, humanized HuNSG-SGM3 and humanized BLT mice were named as NSG-PDX, HuSGM3-PDX and BLT-PDX models, respectively. RNA-seq based mRNA analysis of (**a**) the indicated lymphocyte, (**b**) myeloid cell, and (**c**) NK cell markers genes in NSG-PDX, HuSGM3-PDX and BLT-PDX tumors. Expression is normalized using the expression in matched primary patient derived tumor. (**d**) IF for human CD3, CD8, CD14 and CD163 evaluating tumor immune infiltrates in the indicated PDX tumor models. Cell nuclei were counterstained with DAPI (blue). Scale bars, 20 µm.

### Analysis of Complete HIS-PDX

Having optimized the incorporation of connective tissues, vascular, and immune cells, we combined all aspects of the model to create five HIS-PDX models (3 from solid tumors, 2 from ascites-derived cells) by co-engraftment of primary tumor cells, CA-MSC and EC (iHUVEC + iHDMEC) in BLT humanized mice. IF analysis of the generated HIS-PDX with anti-human mitochondrial antigen confirmed a general humanization of the stroma (Fig. 4a). Compared to a cell line xenograft and traditional PDX, the HIS-PDX model demonstrated increased desmoplasia and a histology more representative of the primary human tumor (Fig. 4b). RNA-seq analysis of solid tumor derived PDX, with a focus on host genes, indicated that HIS-PDX, compared to standard PDX, were more reflective of the gene expression in primary human tumors (Fig. 4c). Comparison across all genes demonstrated a 0.93-0.96 correlation between primary human tumor and HIS-PDX (Fig. 4d).

**Figure 4:**
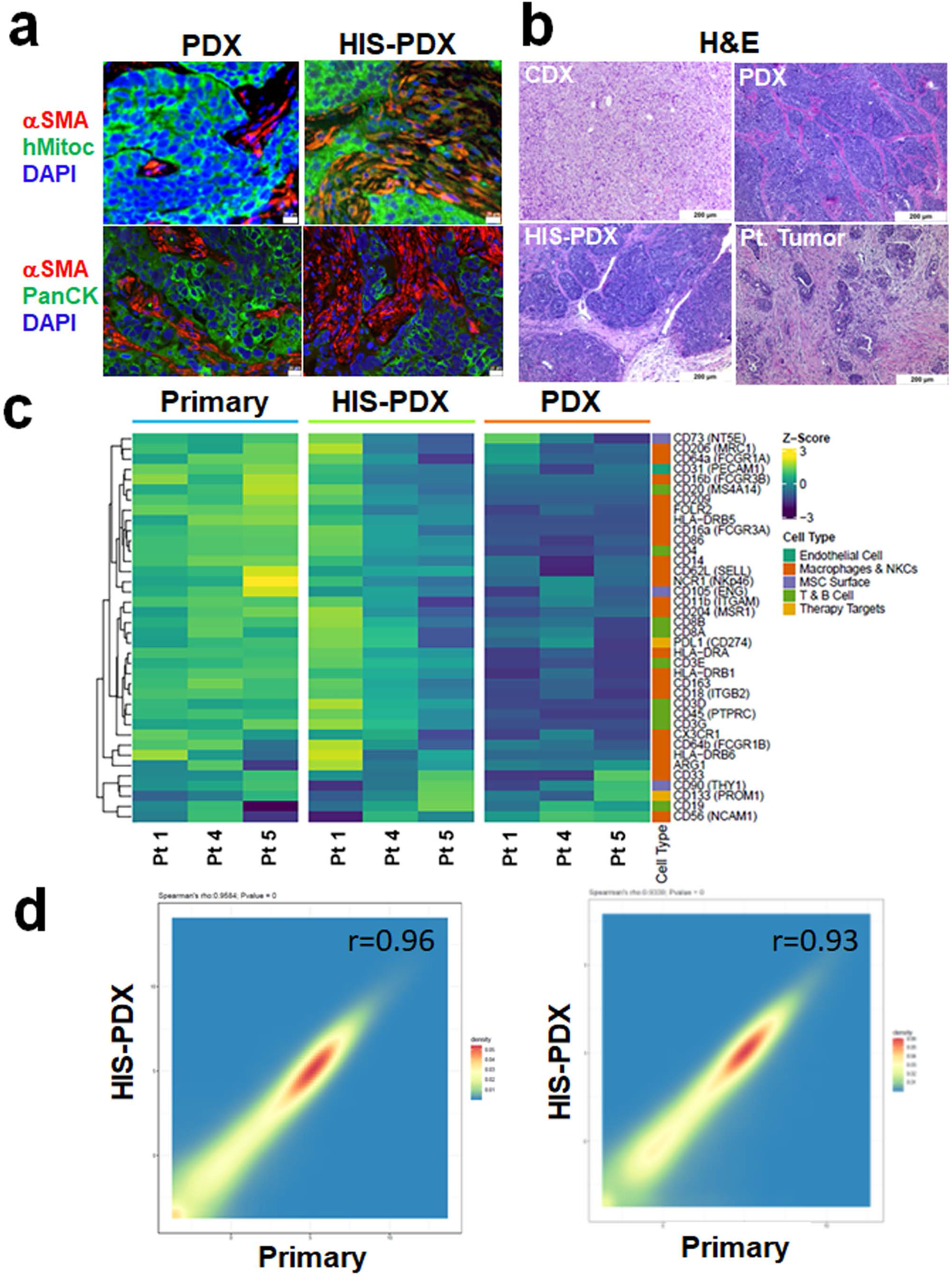
Analysis of complete humanized immune stroma PDX (HIS-PDX). HIS-PDX were created by co-engraftment of PDX TC, CA-MSC and EC (iHUVEC + iHDMEC). (**a**) IF for alpha smooth muscle actin (αSMA; red), and human-specific mitochondrial antigen (hMitoc; green) or pan-cytokeratin (PanCK; green) to label tumor cells in standard and HIS-PDX. Cell nuclei are counterstained with DAPI (blue). (**b**) Hematoxylin and eosin (H&E) staining for a cell line derived xenografts, standard PDX, humanized immune stroma PDX (HIS-PDX) and matched patient’s primary tumor. (**c**) RNA-seq derived heat map for stromal and immune gene expression in solid tumor derived standard PDXs (StndPDX), HIS-PDX, and patient primary tumor. (**d**) Total gene expression correlation between primary human tumor and HIS-PDX. Scale bars in (**a**), 20 µm; scale bars in (**b**), 200 µm.

### Using a HIS-PDX model for therapeutic studies

To test the impact of the humanized stroma on immunotherapy, we first used an adoptive T cell therapy. This model assures specific anti-tumor T cell activity. We used previously validated follicle-stimulating hormone receptor (FSHR)–mediated CAR T-cells(32). As our PDX do not express high levels of the FSHR, we used OVCAR4 cells with confirmed FSHR expression (Supplemental Fig. 3a) combined with the humanized stroma. Tumors were generated in both BLT mice and HuNSG mice. When tumors reached ∼100 mm^3^ we initiated treatment with either mock transfected T cells or FSH-ER transfected T cells (CAR-T cells). Treatment with mock transfected control T cells had no impact on tumor growth in either HuNSG or BLT models (Fig 5a – curves combined). In contrast, CAR-T cell treatment in HuNSG mice resulted in tumor regression. Interestingly, suggesting an increased immunosuppressive TME in the BLT mice, CAR-T cell therapy in the BLT mice resulted in stable disease, but not disease regression.

**Figure 5:**
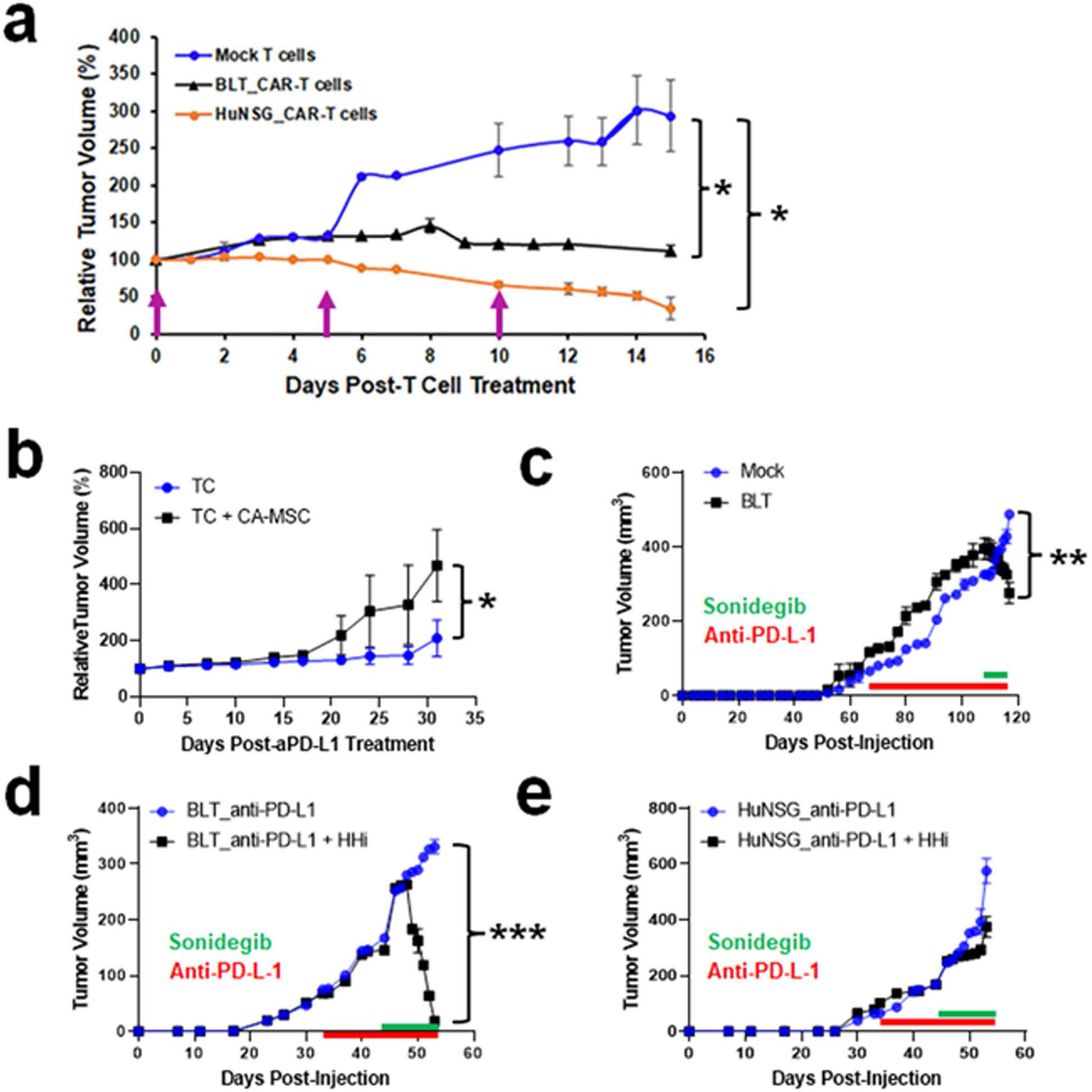
Using a HIS-PDX model for immunotherapeutic studies. (**a**) Impact of humanized stroma on CAR-T cell-based therapy. Arrows indicate days when 4 × 10^6^ mock transfected T cells or CAR-T cells were intraperitoneally injected. (**b**) Tumor growth curves for PDX grown with and without CA-MSC and treated with anti-PD-L1. (**c**) Tumor growth curves of Mock transplanted (non-humanized immune compartment) and BLT humanized PDX treated with anti-PD-L-1 (red bar) in combination with the HHi sonidegib (green bar). (**d**) Tumor growth curves for HIS-PDX in BLT mice treated with anti-PD-L1 in combination with HHi. (**e**) Tumor growth curves of HS-PDX grown in CD34^+^ HuNSG mice and treated with anti-PD-L1 in combination with HHi. For tumor treatment studies above, intraperitoneal injections of avelumab (anti-PD-L1; 10 mg/kg, twice a week) started once PDX tumors reached ∼100 mm^3^ and tumor volumes were monitored twice a week. HHi sonidegib was administered via oral gavage, 10 mg/kg/day qd for 9 days. \**P* < 0.05. \*\**P* < 0.01; \*\*\**P* < 0.001.

While CAR-T cell therapy is in clinical trial for ovarian cancer, it is not currently a clinically available therapeutic strategy. To address strategies that are in the clinic, we evaluated immune checkpoint inhibitor approaches. Prior work has indicated that PD-L1 expressing tumors can respond to PD-1/PD-L1 blockade therapy in HuNSG mice(33). However, response is near universal, which does not reflect the ∼20% response rate in humans(34). We therefore used an HIS-PDX model derived from a patient with immune checkpoint inhibitor resistant disease, to evaluate therapeutic response to anti-PD-L1 (aPD-L1) therapy. RNA-seq analysis and qRT-PCR confirmed that PD-L1 was expressed at increased levels in HIS-PDX models (Supplemental Fig. 3). We recently reported a role for CA-MSC in suppressing response to immune checkpoint inhibitor (ICI) therapy in mice(8). To determine whether CA-MSC impact aPD-L1 therapy in the HIS-PDX, we injected PDX tumor cells with or without CA-MSC into humanized BLTS mice. Tumors were allowed to grow until reaching ∼100 mm^3^ before being treated with intraperitoneal injections of avelumab (aPD-L1; 10 mg/kg, twice a week). Consistent with prior studies, in the absence of human stroma, in BLTS mice with human immune cells avelumab restricted tumor growth (Fig. 5b). In contrast, in the presence of CA-MSC, ∼2 weeks after initiation of therapy clear tumor progression was noted (Fig. 5b). This suggest CA-MSC, directly or indirectly, contribute to tumor immune suppression.

Activation of the hedgehog (HH) pathway is linked to ICI resistance(35,36). Indeed, our group and others have reported that HH signaling pathway inhibitors (HHi) can overcome ICI resistance in mice (8,37). We therefore next tested whether inhibition of HH pathway could increase response to aPD-L1 therapy in the context of a human TME using the HIS-PDX model. HIS-PDX tumor cells were generated in BLT mice. As a control for activity on tumor cells, HS-PDX were generated in mock-transplanted NSG mice. After PDX tumors reached ∼100 mm^3^, we treated mice with avelumab as described above. PDX in both BLT and mock transplanted mice demonstrated clear progression with similar growth kinetics (Fig. 5c). After 6 weeks of avelumab therapy, with clear evidence of therapeutic resistance, we initiated daily treatment with the HHi sonidegib (oral gavage, 10 mg/kg/day qd). Within days of HHi treatment, the HIS-PDX demonstrated profound tumor regression, while HS-PDX in the mock transplanted mice continued to grow (Fig. 5c). Thus, HHi response was dependent upon the human immune compartment.

To better understand the role of the human immune compartment, we repeated studies comparing response in BLT mice (which have an improved myeloid cell compartment) and CD34^+^ HuNSG mice (which have human lymphocytes but generally lack human myeloid cells). Once again, avelumab alone had no effects on the HIS-PDX tumor growth in either the CD34^+^ HuNSG mice or the BLT mice (Fig. 5d, e). Once again, addition of the HHi in the BLT HIS-PDX induced a rapid and profound tumor regression (Fig. 5d). In contrast, addition of HHi therapy in avelumab treated HIS-PDX in the HuNSG mice, had little or no effect (Fig. 5e). Collectively, this data indicates that the human tumor stroma in the HIS-PDX suppresses response to ICI therapy, and that components of the myeloid compartment are essential for tumor response to ICI/HHi therapy.

### HHi treatment is associated with M1 macrophage conversion

To better understand the impact of HHi and the role of the myeloid cell compartment in response to ICI therapy, we performed spatial transcriptomics profiling of ∼1800 human genes on tumor sections of aPD-L1 and aPD-L1/HHi treated HIS-PDX in BLT mice using the digital spatial profiling (DSP) platform (Nanostring Inc.). Evaluation of non-tumor cells (cytokeratin negative) demonstrated that addition of HHi was associated with a significant increase in: (i) cytotoxic immune effectors including perforin-1 (PRF1), granzyme-B, and granulysin-Y, (ii) lymphocyte chemoattractants CXCL9 and CXCL10 (T cell), and CXCL13 (B cell), (iii) an increase in numerous components of antigen presentation, including HLA-DRA, HLA-DRB (MHC-II components), and CTSS (cathepsin-S) (Fig. 6a; FDR-adjusted p<0.05). In line with immune activation, GO-biologic process analyses of differentially expressed genes demonstrated significant enrichment of immune system pathways including TCR signaling, MHC-II antigen presentation, and interferon gamma signaling (Fig. 6b). These findings are consistent with prior studies showing that HHi can promote M2 to M1-reprogramming of TAMs. Indeed, IF analysis of tumors confirmed a significant increase in the expression of HLA-DR^+^ (a marker for M1 TAM)(38) in HHi/aPD-L1 treated tumors vs. aPD-L1 treated tumors (Fig. 6c).

**Figure 6:**
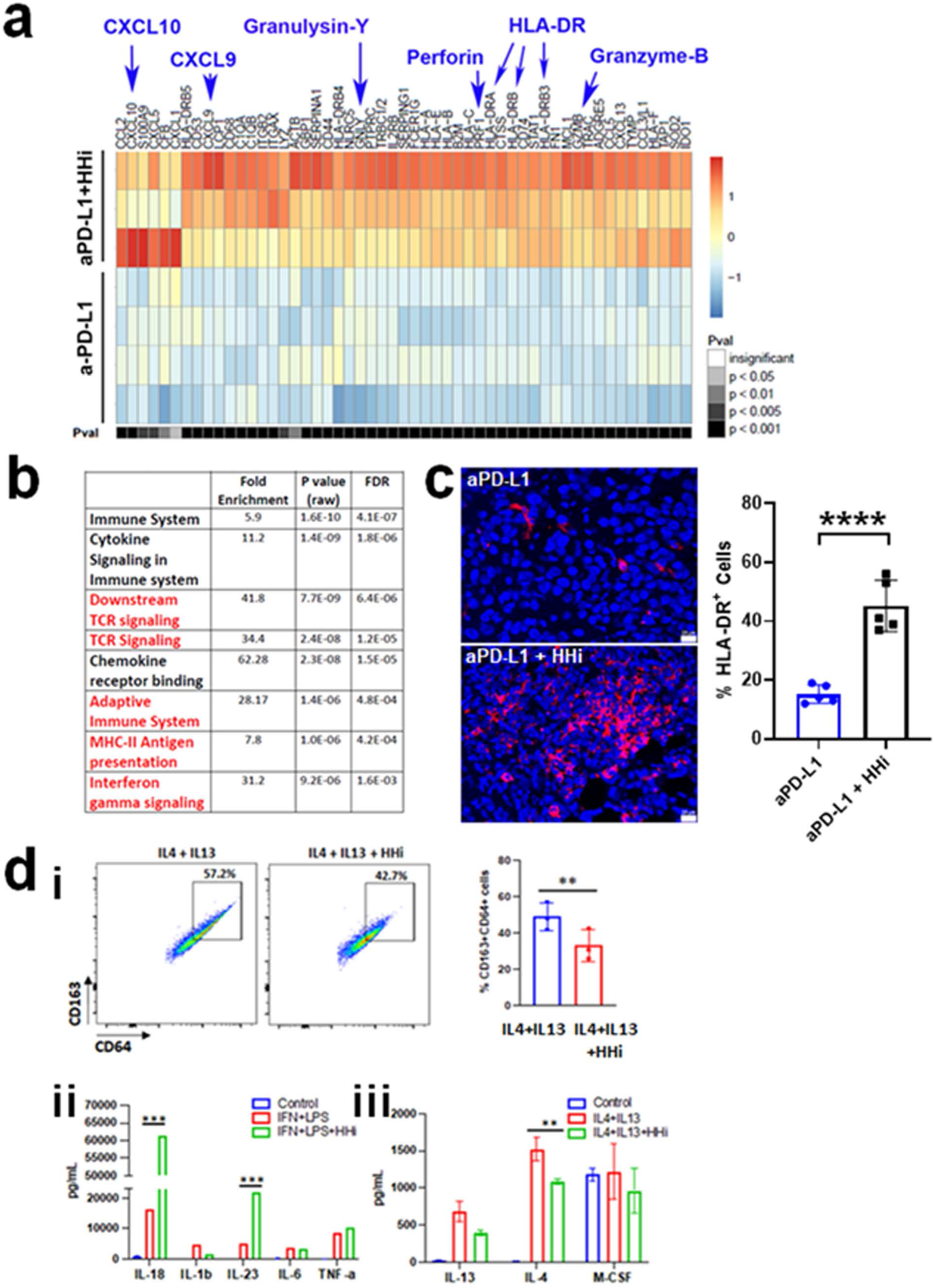
HHi treatment is associated with M1 macrophage conversion. (**a**) Heat map of differentially expressed genes in the cytokeratin negative (non-tumor cell) compartment of HIS-PDX tumors in BLT mice treated in Fig. 5d above measured by DSP GeoMx. Seven regions of interest (ROIs) are shown, including three from samples treated by aPD-L1+HHi, and four from samples treated by a-PDL1. Color bar represents row z-score of log2-transformed and normalized gene expression. P-values shown are after adjusting for multiple comparisons by Benjamini-Hochberg Procedure. FC = fold change. (**b**) GO-biologic process analyses of differentially expressed genes. (**c**) IF of and quantification human HLA-DR^+^ (red) cells in tumors treated with anti-PDL-1 without and with HHi. HLA-DR stain was quantified in 5 high-powered fields from three tumors. (**di**) Flow cytometry analysis and quantification of the abundance of CD64^+^CD163^+^ expression in PBMC-isolated CD14^+^ cells differentiated and polarized toward M2 macrophages (IL4+IL13) in the presence or absence of HHi from 3 replicate experiments. (**dii & diii**) Meso Scale Discovery cytokine expression of the indicated factors in M1 differentiated control and HHi treated macrophages (**dii**) and M2 differentiated control and HHi treated macrophages (**diii**). Each experiment was repeated three times. Samples were run in duplicates. Limma regression models were used in (b). \**p* < 0.05, \*\**p* < 0.01, \*\*\**p* < 0.001, \*\*\*\**p* < 0.0001. Scale bars, 20 µm.

To determine if this is a direct response of TAMs to HHi, we isolated CD14^+^ monocytes from peripheral blood mononuclear cells (PBMC) and primed them toward M2 macrophages using IL4 + IL13 stimulation in the presence or absence of HHi. Consistent with HHi driven M2 to M1 conversion, HHi therapy was associated with a significant reduction in CD163^+^/CD64^+^ macrophages (Fig. 6di). To explore this phenotype change further, we polarized PBMC derived macrophages towards either M1 type macrophages using IFNγ + LPS stimulation, or M2 macrophages using IL4 + IL13 stimulation with or without HHi and evaluated chemokine production. Unprimed macrophages were used as control. The Meso Scale Discovery platform was used to determine cytokine expression levels from macrophage culture supernatants. HHi significantly enhanced the expression of Th1 response cytokines IL-18 and IL-23 in M1 polarized macrophages and decreased the expression of IL-4 in M2 polarized macrophages (Fig. 6d ii-iii).

## Discussion

Greater than 90% of oncology drugs fail in human clinical trials. It has been suggested that a lack of appropriate preclinical models which reflect human disease contributes to this failure (39,40). In this study, we created a novel HIS-PDX preclinical model of human cancer. In contrast to current preclinical models, which lack tumor stroma, this model incorporates human tumor connective tissues, tumor blood vessels, and immune cells. Molecular and protein profiling, and histologic characterization demonstrated that our HIS-PDX model closely reflects that seen in patient primary tumors. As such, we find that this model appears to reflect patient response to both chemotherapy and immunotherapy. Furthermore, this model can be used to explore the immune-suppressive biology of the tumor microenvironment.

The HS-PDX and HIS-PDX models presented here have many strengths compared to other models. These include presence of human stromal and vascular cells, and in the case of the HIS-PDX human thymic education of T cells, and strong NK and myeloid cell compartments. While studies are limited, both models also appear to reflect patient response to chemotherapy and immunotherapy respectively. The HS-PDX model has the strength of being relatively easy to generate and incorporate human stroma, but has the weakness of lacking human immune cells. The HIS-PDX model have the weaknesses that generating BLTS mice is costly and like most humanized immune mouse models, the model has the weakness of having an allogeneic immune system.

Despite an allogeneic immune system of the HIS-PDX model, tumors engraft and grow relatively rapidly. This allows the ability to (i) study mechanisms of immune suppression, and evaluate immunotherapeutics in the context of a near completely human TME. Future studies could combine this model with patient peripheral blood derived hematopoietic stem cells and MSC. This is reported to allow the generation of a humanized mouse with autologous tumors and immune cells (41,42). This combined with patient derived CA-MSC and tumor derived endothelial progenitor cells could result in a completely isogenic humanized model.

Another important feature of the HIS-PDX model is the ability to maintain human tumor stroma. The tumor stroma plays a critical role in promoting tumor growth, metastasis, therapeutic resistance and in regulating anti-tumor immunity (43,44). We demonstrated the presence of many human stromal cell types including CAF, EC, TILs and TAMs in our HIS-PDX model (Figs 1-4). As CA-MSC can be easily isolated from primary tumors and expanded for several months ex-vivo without losing their phenotype (16,17), they can be used as a source of stromal cells even in later passage PDX and in large scale therapeutic studies. Endothelial cells can be generated in the model with either primary tumor derived CD34^+^KDR^+^NRP1^+^cells or immortalized endothelial cell lines. The primary human tumors cells have the advantage of being autologous with tumor cells and in generally provided a more robust human vasculature. However, their numbers are more limited. Immortalized endothelial cell lines offer a significant advantage of ease of use and easy scalability.

In our studies we found the NSG-BLT/S model provides a superior human immune system compared with the Hu-NSG and Hu-NSG-SMG3 models. Both NK cells and myeloid cell compartments were well represented in the BLT model and generally lacking in the Hu-NSG model. We also note that, in our hands, Hu-NSG-SGM3 mice had significant issues with graft vs. host disease that was not seen in the BLT mice.

Importantly, the HIS-PDX model can be used as a platform to identify compounds to improve human immunotherapeutic response and using different humanize immune models to study the immune cell compartments related to this response. In the HIS-PDX model, consistent with clinical trial outcomes for ovarian cancer patients (45), ICI therapy alone had no effect on tumor growth. However, in line with numerous recent studies, we found HHi could overcome the stromal immune suppression. These models also highlight and represent a novel tool to study the complex role of human myeloid cells in regulating antitumor immunity. In the HIS-PDX in HuNSG mice, which lack a human myeloid cell compartment, CAR-T cell therapy induced tumor regression while the same treatment in BLT mice resulted in only stable disease. This is consistent with a well-documented role of myeloid cells in immune suppression. In parallel, an immune suppressive role for the myeloid compartment was observed with single ICI therapy; ICI therapy demonstrated disease control in HIS-PDX in HuNSG mice, but little or no effect in BLT mice. This is consistent with a reported immunosuppressive role of CA-MSC, via recruitment and polarization of M2 TAMs, driving the resistance to ICI therapy (8). However, at least in the HIS-PDX model, the myeloid cell compartment is also important for tumor mediated immune rejection; tumor immune rejection in response to dual ICI/HHi therapy was not observed in Hu-NSG but was observed in the BLT mice.

In conclusion, we created HS-PDX and HIS-PDX models by co-transplanting three different types of human cells (TC, CA-MSC and EC) into NSG or humanized BLT mice. These model, by faithfully recapitulating human tumor stroma, serves as a better tool for therapeutic discovery and development and could revolutionize preclinical drug testing.

## Methods

### Human tumor tissues and tumor cell isolation

All human tumor tissues were collected with IRB approval (PRO17090380) by the University of Pittsburgh. Tumor tissues were obtained via the Magee-Womens Hospital and the University of Pittsburgh Biospecimen Core (http://www.pittbiospecimencore.pitt.edu/). All participants provided a written consent for tumor tissue collection. Tumor tissues of high-grade serous ovarian carcinoma (HGSOC) confirmed by pathologists were used for this study. Human tumor cells were isolated from primary human omental metastases with human tumor cell isolation kit (Miltenyi Biotec, Cat# 130-108-339) using MACS^®^ cell separation technology following the manufacturer’s instructions. Tumor cells were freshly used for establishing patient-derived tumor xenografts (PDXs) or cryopreserved until use. Alternatively, tumor cells were isolated from HGSOC associated ascites.

### Cell culture

Cancer-associated mesenchymal stem cells (CA-MSC) were isolated from ovarian tumor tissues and cultured as previously described(17).

Human induced pluripotent stem cell (iPSC) line was purchased from ALSTEM, LLC (Richmond, CA 94806; Cat#: iPS12). This cell line was generated by transiently introducing episomal plasmids encoding the human transcription factors into human MSC-derived from an adult bone marrow. Human iPSCs were cultured under feeder-free conditions according to procedures outlined by the manufacturer (ALSTEM) except for dissociation of iPSCs. We found that the EDTA (0.5 mM in Dulbecco’s phosphate-buffered saline (DPBS) for ∼5 -10 min)-dissociated iPSC clusters (aggregates) can be successfully used for passaging and cryopreservation of iPSCs, consistent with the findings by other groups(46).

Differentiation of EPC from iPSCs was performed as previously described(47). Briefly, human iPSCs grown on a Matrigel-coated surface in mTeSR1 (STEMCELL Technologies) were dissociated into single cells with Accutase (Life Technologies) at 37°C for 5 min and then seeded onto a Matrigel-coated cell culture dish at 50,000 cell/cm^2^ in mTeSR1 supplemented with 5 µM ROCK inhibitor Y-27632 (Sigma, Cat#: Y0503) (day-3) for 24 hr. Cells were then cultured in mTeSR1, changed daily. At day 0, cells were treated with 6 µM CHIR99021 (Sigma, Cat#: SML1046) for 2 days in LaSR basal medium consisting of advanced DMEM/F12 (Fisher Scientific, Cat#: 12-634-010 (500 mL) or 12-634-028 (12 × 500 mL), 2.5 mM GlutaMAX (Fisher Scientific, Cat#: 35-050-061), and 60–100 µg/ml ascorbic acid (Sigma, Cat#: A8960). After 2 days, CHIR99021-containing medium was removed and cells were maintained in LaSR basal medium without CHIR99021 for additional 3-16 days.

Day 5-18 differentiated cells were dissociated with Accutase for 10 min and CD34^+^ population was purified by magnetic-activated cell sorting (MACS) with an EasySep Magnet kit (STEMCELL Technologies) using a CD34 antibody according to the manufacturer’s instructions. The purified CD34^+^ cells were plated on collagen-IV-coated dishes (BD BioCoat) in endothelial cell growth medium (PromoCell GmbH, Fisher Scientific, Cat#: 50-306-195) or EGM-2 medium (Lonza) and split every 3-7 days with Accutase.

Immortalized human dermal microvascular endothelial cell line (iHDMEC) TIME (ATCC^®^ CRL-4025™), and immortalized human umbilical vein endothelial cell line (iHUVEC) HUVEC/TERT 2 (ATCC^®^ CRL-4053) were purchased from ATCC (Manassas, VA, USA). EC were cultured following the manufacturer’s instructions.

### Flow cytometry analysis

Cells were stained with mouse (m) or human (h)-specific antibodies-fluorophores in varying combinations as indicated in figures and figure legends: hCD31 (PECAM-1)-APC, hCD34-FITC, hKDR (VEGFR2)-PE, hCD73-PE, hCD90-FITC,h CD105-APC, hCD45-FITC (clone HI30, BD Cat#: 555482), hCD3-APC (clone HIT3a, BD Cat#: 555342), hCD14-PE (clone 63D3, BioLegend, Cat#: 367104), mCD45-PE (clone 30-F11, BioLegend, Cat# 103106), hCD45-APC-C7 (clone HI30, BioLegend, Cat# 304014), CD163 (clones RM3/1, Biolegend), CD64 (clone 10.1, Biolegend), CD14 (clone M5E2, Biolegend), HLA-DR (clone LN3, eBioscience) and corresponding isotype controls (Supplemental Table 2). Cells were stained in DPBS buffer containing 2% fetal bovine serum (FBS) on ice for 30 minutes, washed, resuspended in fresh DPBS buffer containing 1% FBS and analyzed on a CytoFLEX S flow cytometer (Beckman Coulter). Dead cells were excluded by 4′,6-diamidino-2-phenylindole (DAPI).

We used fluorescence-activated cell sorting (FACS) to purify MSC and EPC as previously described(17,28,47). Cells were stained and washed as above. Sorting was performed at the Magee Womens Research Institute Flow Cytometry Core using a BD FACSAria™ III sorter with FACSDiva software (BD Biosciences).

### Animals

All animal experiments were conducted in accordance with standard regulations and were approved by the University of Pittsburgh Institutional Animal Care and Use Committee. Immunodeficient NOD.Cg-*Prkdc^scid^ Il2rg^tm1Wjl^*/SzJ (NSG) (Strain #:005557; RRID:IMSR_JAX:005557) and NOD.Cg-*Prkdc^scid^ Il2rg^tm1Wjl^* Tg(CMV-IL3,CSF2,KITLG)1Eav/MloySzJ (NSG-SGM3) (Strain #:013062; RRID:IMSR_JAX:013062) were purchased from the Jackson Laboratory (Bar Harbor, ME, USA). Mice were housed in a standard dark/light cycle with ambient temperature and humidity conditions in the University of Pittsburgh animal facility.

NSG mice were used as hosts for establishment of 3 humanized mouse models including humanized NSG mice (HuNSG), BM-liver-thymus (BLT) and BM-liver-thymus-spleen (BLTS) as previously described(13,31). NSG-SGM3 mice were used for establishment of humanized NSG-SGM3 mouse model (HuSGM3) as described(13). Humanized HuSGM3 and a fraction of HuNSG were purchased from the Jackson Laboratory.

### Establishment of PDX models and humanized PDX models

Each of patient-derived xenograft (PDX) models was established by transplanting patient tumor cells (1LJ×LJ10^6^) with or without CA-MSC (0.5 ×LJ10^6^) and/or EC (0.25 ×LJ10^6^ iHDMEC + 0.25 ×LJ10^6^ iHUVEC) into the flank of each of NSG mice. PDX tumor cells were isolated using MACS^®^ cell separation technology and serially passaged into NSG mice. Passages 1–3 PDX tumor cells with or without CA-MSC and/or EC were transplanted into the flank of each of humanized mice to establish humanized PDX models.

Tumor volumes were measured twice weekly by a digital caliper and calculated using the modified ellipsoid formula V = (L × W × W)/2, where V is tumor volume, L is tumor length and W is tumor width. Tumor growth curves were graphed in Prism 9 (GraphPad) or Excel for Microsoft 365 (Microsoft Corporation, Redmond, WA, USA).

### *In vivo* therapeutic studies

Cell line xenografts with CA-MSC derived human stroma were generated as previously described(16,17). Briefly 200K tumor cells and 200K CA-MSC were mixed and injected subcutaneously in mice. Treatment was initiated seven days after inoculation. DBZ 10 μmol/kg was administered IP daily for 21 days. Anti-human IL-6 (AB-206-NA, R&D Systems) 10 mg/kg was administered intraperitoneally (IP) twice weekly for six doses. TG101209 100 mg/kg was administered IP daily for 21 days.

Standard PDX models (NSG-PDX) were established in immunodeficient NSG mice, while humanized PDX models (HS-PDX, HuNSG-PDX, BLT-PDX and BLTS-PDX) were established in NSG or the corresponding humanized mouse models, HuNSG, BLT and BLTS as described above. For carboplatin studies, when tumors reached ∼50mm^3^, they were treated with three weekly doses of carboplatin (75mg/kg) and taxol (20mg/kg) administered IP as previously described (48). For other therapeutic studies, when tumors reached ∼100 mm^3^, mice were randomized before treatment. For FSHR–mediated CAR T-cell therapy, mock transfected T cells or FSH-ER transfected T cells (CAR-T cells) were prepared as previously described(32). Four million mock transfected T cells or CAR-T cells in 100 μl of PBS were intraperitoneally injected at 5-day intervals for a total of 3 doses. For immune checkpoint inhibitor therapy with or without HHi, mice were treated as following: (a) BLTS mice were transplanted with PDX tumor cells with or without CA-MSC and treated with IP of avelumab (anti-PD-L1; 10 mg/kg) twice a week; (b) mock NSG and humanized BLT mice were transplanted with PDX tumor cells with CA-MSC and treated with avelumab as in (a) for 50 days, followed by avelumab (IP) in combination with oral gavage of HHi sonidegib (10 mg/kg/day) daily and for additional 9 days; (c) HuNSG and BLT mice were transplanted with PDX tumor cells with CA-MSC and EC and treated with avelumab as in (a) for 13 days, followed by avelumab (IP) and sonidegib (oral gavage) as in (b) for additional 6 days. At end of experiments, mice were euthanized, and tumor and spleen tissues were collected for RNA isolation, Hematoxylin and Eosin (H&E) staining, IF and/or flow cytometry analysis.

### RNA isolation and RT-qPCR

Total RNA was isolated from patient or PDX tumor tissues using RNeasy-mini or RNeasy-Plus-Micro kits (Qiagen). cDNA was synthesized using SuperScript™ III First-Strand Synthesis System (ThermoFisher Scientific, Cat#: 18080051). Quantitative PCR (qPCR) reactions were performed in triplicate with SYBR Green Supermix (Bio-Rad, Cat#: 1725271). Glyceraldehyde-3-phosphate dehydrogenase (GAPDH) or hypoxanthine guanine phosphoribosyl transferase 1 (HPRT1), was used as an internal control. PCR primer sequences were listed in Supplemental Table 3.

### RNA sequencing

Patient and PDX tumor tissues (passage 1 or passage 2) from 5 different PDX models were used for RNA isolation as described above. The amount of RNA was measured using the Nanodrop. RNA quality (A260/A280 ≥ 2.0, A260/A230 ≥ 2.0) was verified using the Nanodrop and Agilent 2100. RNAs with RNA integrity number ≥ 6.3 RIN (RNA integrity number) and flat base line determined by Agilent 2100 were used for RNA sequencing (RNA-seq) analysis.

After RNA quality control, mRNA was enriched, double-stranded cDNA was synthesized, ends were repaired, poly-A & adaptors were added, fragments were selected, and PCR was performed. cDNA libraries were prepared. The quality of cDNA libraries was assessed. Libraries were sequenced. All of those RNA-seq procedures were performed by Novogene (Sacramento, CA). RNA-seq data analyses were performed by Novogene and Dr. Hui Shen’s research group. Raw reads were first trimmed of adapters with trim_galore and then mapped to GRCm38 primary assembly with STAR v2.7.8a using options ‘--twopassMode Basic’ and ‘--quantMode GeneCounts’ to directly output counts for all features from GENCODE human release 39. Libraries averaged 31 million uniquely mapped fragments and an average base quality of 35. Data were quality controlled using Fastq Screen (https://www.bioinformatics.babraham.ac.uk/projects/fastq_screen/) to check for mouse contamination. There were no statistical differences between PDX groups, with 5% of sequencing reads originating from mouse on average in PDXs (0% in primary tumors). Heatmaps were generated from library-size normalized counts centered across genes (z-scores). Heatmaps were generated using the R package ComplexHeatmap(49).

### H&E staining

Tumor tissues were paraffin embedded. Paraffin-embedded tissues were sectioned at 5 μm thickness. H&E staining was performed at the Pathology Cores for Animal Research at the Magee-Womens Research Institute or Pitt Biospecimen Core at University of Pittsburgh.

### Immunofluorescence (IF)

IF was modified based on a previous publication(50) and an online protocol developed by R&D Systems (https://www.rndsystems.com/resources/protocols/protocol-preparation-and-fluorescent-ihc-staining-paraffin-embedded-tissue). Briefly, the slide-mounted paraffin-embedded tissue sections (5 µm) were deparaffinized in xylene and rehydrated in ethanol and PBS before antigen retrieval in sodium citrate buffer (10 mM, pH 6.0) for 10 min at 95°C. The sections were then permeabilized and blocked with blocking solution (0.3% Triton X-100, 5% normal goat serum and 3% BSA in PBS) for 60 minutes at room temperature. Sections were incubated overnight at 4°C with primary antibodies diluted in 0.05% Triton X-100, 0.8% normal goat serum and 0.5% BSA in PBS. The sections were extensively washed 3 times with PBS, then incubated at room temperature for 60 minutes with secondary antibodies diluted in 0.05% Triton X-100, 0.8% normal goat serum and 0.5% BSA in PBS to a final dilution of 1:500. Finally, the sections were washed with PBS, and mounted in antifade mounting medium with DAPI. Digital images were viewed with the Leica DM4 B upright microscope (Leica Chicago, IL) and collected with allied Leica LAS X 3.3 Software. For negative controls, primary antibodies were omitted. Primary antibodies and secondary antibodies for IHC are listed in Supplemental Table 3. Quantification of stain was done in a minimum of 5 high-powered fields for 3 tumors for each treatment group and compared via ANOVA.

Digital image data were quantified by ImageJ (http://imagej.nih.gov/ij/docs/index.html). Figure panels were composed with Photoshop software (Adobe Systems, Mountain View, CA, USA) or PowerPoint for Microsoft 365 (Microsoft Corporation, Redmond, WA, USA) for archival purposes.

### Peripheral blood monocyte isolation and macrophage differentiation

Peripheral blood mononuclear cells (PBMC) were freshly isolated by density-gradient centrifugation using Ficoll Paque Plus (Sigma-Aldrich) for 50 minutes at 400 × g. Deidentified human Buffy Coat samples were purchased from Vitalant fulfilling the basic exempt criteria 45 CFR 46.101(b) in accordance with the University of Pittsburgh IRB guidelines. Monocytes were then isolated with CD14^+^ microbeads (MACS Miltenyi) and incubated for 6 days in RPMI/10%FBS and 1% penicillin/streptomycin solution (Sigma) supplemented with 50 ng/mL human GM-CSF (R&D Systems) to stimulate macrophage differentiation. Once differentiated, macrophages were washed and polarized toward M1 or M2 phenotypes by incubation with 40 ng/mL LPS (Sigma-Aldrich) and 100 ng/mL IFNγ (R&D Systems) or 50 ng/mL and IL4 and IL13 (R&D Systems) for 48 hours, respectively. Differentiated macrophages cultured with RPMI alone were used as control. In certain experiments, 10 µM hedgehog inhibitor (Sorafenib) was added to the priming conditions. After treatments, macrophages were washed and kept in culture in fresh media for an additional 24 hours. Then, cells were collected and analyzed by flow cytometry whereas supernatants were stored at -80°C for subsequent analysis by Meso Scale Discovery as indicated below.

### Cytokine multiplex immunoassay

Secreted cytokine levels of human differentiated and polarized macrophages treated or not with hedgehog inhibitor were measured using the U-Plex Human Macrophages M1 Combo (IL-1β, IL-6, IL-12p70, IL-18, IL-23, IP-10, MCP-1, MIP-1α, TNF-α) and Macrophages M2 Combo (Eotaxin-2, IL-4, IL-10, IL-13, M-CSF, MDC, TARC) kits from Meso Scale Discovery according to the manufacturer’s instructions. All MSD assays were read using a Meso QuickPlex SQ 120 and analyzed using Discovery Workbench (MSD).

### Digital spatial profiling (DSP) library preparation and sequencing

GeoMx^TM^ DSP was used to spatially quantify the intensity of ∼1800 transcripts simultaneously including known immune markers and cancer driver genes (GeoMx® Cancer Transcriptome Atlas, CTA). Formalin-fixed paraffin-embedded tissues (FFPE) sections (5 µm) prepared from HIS-PDX tumors were deparaffinized and subjected to antigen retrieval procedures. The sections were then incubated overnight with a cocktail of DSP antibodies (probes) tagged with unique photocleavable DNA oligos. After staining, the slides were scanned using the GeoMx DSP instrument (NanoString Technologies) to produce digital fluorescent images of each section. Regions of interest (ROIs) were generated and segmented into defined tissue compartments by epithelial marker pan-cytokeratin (PanCK), immune cell marker CD45 and non-epithelial compartment (stroma). DNA oligos from these compartments were released, collected, dispensed into a 96-well plate, hybridized to optical barcodes. Libraries were sequenced on an NextSeq 500 (Illumina Inc.) instrument at Health Sciences Sequencing Core at UPMC Children’s Hospital of Pittsburgh. Two hundred million paired-end reads were generated.

### DSP data analysis

Sequencing files were converted from FastQ to Digital Count Conversion (DCC) format using The GeoMx NGS Pipeline (v1.0). Reads were processed to clip adaptors, merge overlapping mates, align to the Readout Tag Sequence-ID (RTS-ID) barcodes, and remove PCR duplicates by Unique Molecular Identifiers (UMI). The resulting read count matrix was processed in R (v4.2.1) for further analysis. Data were passed through robust QC procedure to remove ROIs of low nuclei count, low surface area or binding density, low quality probes, high background noise, or identified as global or local outliers. The remaining data were upper quartile normalized and log2-transformed. Genes differentially expressed between treatment groups were detected by *limma* (v3.52.2) regression models and filtered by FDR-adjusted *P*<0.05.

### Statistical Analysis

Data are presented as means ± standard deviation (SD) and were analyzed using Graphpad Prism version 9 (GraphPad Software, Inc., San Diego, CA, USA). Statistical significance between two groups was evaluated by two-tailed Student’s *t* test. Statistical significance among three or more groups was evaluated by one-way analysis of variance (ANOVA), followed by Tukey’s multiple comparisons test. *P* < 0.05 was considered statistically significant.

## Acknowledgements

This work was supported by National Cancer Institute Grants 5R01CA211913 and 5R01CA218026.

## Competing Interests

The authors have declared that no competing interest exists.

## Author contributions

D.Y. and R.J.B. conceived and designed the study, analyzed data and wrote the manuscript with inputs from all of the authors. D.Y., H.T., S.C.M., S.C., S.B. and M.T.B. performed experiments. I.B., B.K.J. and H.S. analyzed RNA-Seq data and contributed to the corresponding figures. R.A., R.B. and T.C.B. analyzed DSP data and contributed to the corresponding figure. J.J.P. and J.R.C. provided reagent information and reagents. T.R.S. contributed to histologic evaluation of tissue and manuscript review. R.J.B. supervised the project.

## Supplemental Figure Legends

**Supplemental Figure 1: Related to Figures 1&2. CA-MSC increase tumor metastasis to ovaries.** Subcutaneous co-engraftment of ovarian patient-derived xenograft (PDX) tumor cells and cancer-associated mesenchymal stem cells (CA-MSC) increases tumor micrometastasis to ovaries as demonstrated by fluorescent IHC analysis of human cytokeratin (CK; green). Cell nuclei were counterstained with 4′,6-diamidino-2-phenylindole (DAPI; blue) Scale bars, 20 µm.

**Supplemental Figure 2: Related to Figure 2. Using iPSC-EPC to generate human endothelial cells.** Using human induced pluripotent stem cells (iPSC)-derived endothelial progenitor cells (EPC) to make blood vessels. (**a**) Flow cytometry analysis of CD34 and CD31 expression in unsorted and sorted CD34^+^ EPC. For differentiation of human iPSC to EPC, human iPSC were cultured on Matrigel-coated dishes in LaSR basal medium containing 6 µM CHIR99021 for 2 days followed by additional 10 days in LaSR basal medium. CD34^+^ cells were enriched by magnetic-activated cell sorting (MACS). (**b**) **(i)** Caov3 cell line xenografts with or without CA-MSC and iPSC-EPC. No tumor was formed when engrafted with CA-MSC and iPSC-EPC, but without Caov3. **(ii)** PDX with CA-MSC without or without iPSC-EPC. (**c**) IF evaluation of human tumor vascular antigens in CDX with iPSC-EPC. CD31 (red) and kinase insert domain receptor (KDR; also known as vascular endothelial growth factor receptor 2) were used as general vascular markers. Cell nuclei were counterstained with DAPI (blue). Scale bars, 40 µm.

**Supplemental Figure 3: (a)** RT-qPCR of expression of human FSHR in the indicated ovarian cancer cell lines. Data was normalized to a housekeeping gene, GAPDH, then normalized to OVCAR3. The identity of FSHR RT-PCR product was confirmed by sequencing. (**b**) Fold change of CD274 (PD-L1) gene expression in 5 different humanized immune stroma PDX (HIS-PDX) models vs. 5 different standard PDX (PDX) models. (**c**) RT-qPCR analysis of CD274 (PD-L1) gene expression in standard PDX and HIS PDX models. GAPDH was used as an internal control. (**c**) IF analysis of CD274 (PD-L1) (red) protein expression in standard PDX and HIS PDX models. Cell nuclei were counterstained with DAPI (blue). Scale bars, 20 µm.

## Notes

### Competing Interest Statement

The authors have declared no competing interest.

